# When do opposites attract? A model uncovering the evolution of disassortative mating

**DOI:** 10.1101/2020.05.19.104190

**Authors:** Ludovic Claude Maisonneuve, Thomas Beneteau, Mathieu Joron, Charline Smadi, Violaine Llaurens

## Abstract

Disassortative mating is a rare form of mate preference that promotes the persistence of polymorphism. While the evolution of assortative mating, and its consequences on trait variation and speciation have been extensively studied, the conditions enabling the evolution of disassortative mating are still poorly understood. Mate preferences increase the risk of missing mating opportunities, a cost that can be compensated by a greater fitness of offspring. Heterozygote advantage should therefore promote the evolution of disassortative mating, which maximizes the number of heterozygous offspring. From the analysis of a two-locus diploid model, with one locus controlling the mating cue under viability selection and the other locus coding for the level of disassortative preference, we show that heterozygote advantage and negative frequency-dependent viability selection acting at the cue locus promote the fixation of disassortative preferences. The conditions predicted to enable the evolution of disassortative mating in our model match the selection regimes acting on traits subject to disassortative mating behavior in the wild. In sharp contrast with the evolution of assortative preferences, we also show that disassortative mating generates a negative frequency-dependent sexual selection, which in turn disadvantages heterozygotes at the cue locus, limiting the evolution of disassortative preferences. This negative feedback loop could explain why this behavior is rare in natural populations.

## Introduction

The evolution of mate preferences is puzzling because preferences increase the risk of missing mating opportunities, which may incur significant fitness costs. While the evolution of assortative mating has been reported in many species, disassortative mating is more scarcely observed (Janicke et al., 2019; Jiang et al., 2013), suggesting that the ecological conditions enabling its evolution could be more restrictive. Here we build a general approach aiming at investigating the selection regimes allowing the evolution of disassortative mating using a mathematical model.

The multiple costs associated with mate choice tend to generate direct selection against the evolution of mate preferences (see (Pomiankowski, 1987) for a review), and may further limit the evolution of disassortative mating (see (Kopp and Hermisson, 2008; Otto et al., 2008; Pomiankowski, 1987; Schneider and Bürger, 2006) for theoretical studies). These costs of choosiness are generally separated into fixed and relative costs (Otto et al., 2008). Relative costs depend on the distribution of the mating cue within population. For example, relative costs of choosiness may emerge from the increased investment in mate searching, because an individual needs to investigate several mates to find a suitable one. Increased sampling effort can be costly in time Kruijt and Hogan (1967), in energy (as empirically estimated in antilopes Byers et al. (2005)) and may enhance predation risk, for instance in patrolling animals Hughes et al. (2012). Evaluation effort increases with the proportion of unpreferred males, implying growing relative costs of choosiness when the preferred cue is rarely displayed in the population. In addition, mate rejection by choosy individuals can also incur relative fitness costs, as in the case of males harassment: in the fly species *Musca domestica*, males jump on females’ back to initiate mating and choosy females have to kick unpreferred males to avoid mating (Sacca, 1964). The number of males to kick out decreases with the proportion of preferred males. By contrast, fixed costs associated with mate choice do not depend on the composition of the population. For instance, metabolic costs may emerge from the mechanisms underlying mate choice, requiring specialized morphological, physiological and cognitive changes (see Rosenthal (2017) for a review). For example, in the self-incompatibility system in the genus *Brassica*, mate choice involves a specialized receptor-ligand association (Hiscock and McInnis, 2003), so that the evolution of self-incompatibility is associated with metabolic costs induced by the production of the specific proteins.

Despite these costs, mate choice is ubiquitous in nature (Backwell and Passmore, 1996; Barrett, 1990; Cisar, 1999; Hiscock and McInnis, 2003; Jiggins et al., 2001; Merrill et al., 2014; Savolainen et al., 2006) indicating that mate preference evolves readily and that choosy individuals enjoy benefits compensating those costs. Choosy individuals may enjoy direct benefits (Wagner, 2011) (for instance through beneficial sexually transmitted microbes (Smith and Mueller, 2015), or by decreasing risk of pre-copulatory cannibalism (Pruitt and Riechert, 2009)), as well as indirect benefits associated with mate preferences through an enhanced quality of their offspring (Byers and Waits, 2006; Drickamer et al., 2000; Jiggins et al., 2001; Petrie, 1994; Sheldon et al., 1997; Welch et al., 1998).

Viability selection acting on mating cues, by generating indirect selection on preferences, may thus promote their evolution (Fisher, 1930). Such indirect selection is caused by genetic associations between mating preference and mating cues (linkage disequilibirum) (Barton and Turelli, 1991; Ewens, 1979; Kirkpatrick and Ravigné, 2002), generated during zygote formation because of mate preferences. The indirect effect of viability selection, that acts directly on mating cues, on the evolution of mate preferences, first identified by Fisher, has now been confirmed in many theoretical studies (Barton and Turelli, 1991; Heisler, 1984; O’Donald, 1980a). Preference based on a selectively neutral mating cue may also evolve if the cue is correlated with an adaptive trait due to linkage disequilibrium between preference and an adaptive trait (Heisler, 1985). A growing number of empirical evidence showing that female choice does improve offspring fitness is reported (Byers and Waits, 2006; Drickamer et al., 2000; Petrie, 1994; Sheldon et al., 1997; Welch et al., 1998), suggesting that preferences generate linkage disequilibria between preference alleles and other combinations of alleles favored by viability selection. The indirect selection may thus be a major driver of the evolution of mate choice.

Once mate preferences are established in the population, they generate sexual selection on the traits exhibited by individuals during courtship, that may drive the evolution of extravagant traits in males, following a Fisherian runaway (Fisher, 1930; Gomulkiewicz and Hastings, 1990; Greenspoon and Otto, 2009; Kirkpatrick, 1982; Lande, 1981; O’Donald, 1980b; Otto, 1991; Veller et al., 2020). The evolution of mate preferences thus involves complex evolutionary processes where preferences co-evolve with the cues displayed by the chosen individuals. This co-evolution has been observed in natural populations (Grace and Shaw, 2011; Higginson et al., 2012) and in experimental studies (Brooks and Couldridge, 1999; Miller and Pitnick, 2002), underpinning the importance of sexual selection feedbacks on the evolution of mate preferences.

The different selection regimes acting on mating cues can therefore drive the evolution of different mating patterns, through indirect selection. Disruptive selection on mating cue, has been demonstrated to promote assortative preferences (Bank et al., 2012; de Cara et al., 2008; Dieckmann, 2004; Gavrilets, 2004; Kirkpatrick, 2000; Otto et al., 2008). By contrast, selection conferring fitness advantages to intermediate phenotypes is often thought to promote disassortative mating (Kirkpatrick and Nuismer, 2004; Kondrashov and Shpak, 1998). Nevertheless, the selection regimes enabling the evolution of disassortative mating are much less studied than the selective pressures involved in the evolution of assortative mating, extensively investigated in the context of speciation (Gavrilets, 2004; Kopp et al., 2018).

Disassortative mating has been documented only in a few cases. The best documented cases are the MHC loci in humans and mice, where females prefer males with a genotype different from their own (Wedekind et al., 1995). MHC genes are involved in specific recognition of pathogens, and host-pathogens interactions classically generate negative frequency dependent selection and/or heterozygote advantage (recognition of a larger range of pathogens) (Piertney and Oliver, 2006). Such balancing selection regimes are thought to promote disassortative mating at MHC loci (Ihara and Feldman, 2003; Penn and Potts, 1999; Slade and McCallum, 1992). Using numerical simulations in a haploid model, Howard and Lively (2003, 2004) confirm that host-pathogens interactions at MHC loci promote the emergence of disassortative mating, although they never observed the fixation of this mating behavior in the population. In a more general model, Nuismer et al. (2008) observe that sexual selection due to non-random mating generates indirect selection on preference that hampers the fixation of disassortative mating in the population. Despite this limitation, the frequency of disassortative mating can be high when viability selection strongly promotes this behavior. In an extension of Nuismer et al. (2008)’s model, Greenspoon and M’Gonigle (2014) show that maternal transmission of pathogens leads to higher levels of disassortative mating because mothers have increased fitness when they produce offsprings with MHC genotypes different from their own, that might be more effective in eliminating transmitted pathogens.

Other cases of disassortative mating in traits unlinked to immune functions have been reported, such as disassortative mating based on the plumage coloration in the white throated sparrow (Throneycroft, 1975), or on the wing color pattern in the mimetic butterfly *Heliconius numata* (Chouteau et al., 2017). In both cases, one cue allele is linked to a genetic load (Jay et al., 2019; Tuttle et al., 2016), so that disassortative mating may increase offspring fitness through an increased viability of heterozygotes. In both cases, cue alleles associated with a genetic load are dominant to other alleles, suggesting that dominance among cue alleles may play a role in the evolution of disassortative mating. Numerical simulations designed from the specific case of *Heliconius numata*, confirm that heterozygote advantage at the locus controlling color pattern variation may promote the emergence of disassortative mating (Maisonneuve et al., 2019).

Other theoretical studies have focused on the effect of disassortative mating on the persistence of variations at the cue locus, illustrating that this mate preference may limit the purging of maladaptive cue alleles, and therefore promotes higher levels of polymorphism at the cue locus (Falk and Li, 1969; Ihara and Feldman, 2003; Karlin and Feldman, 1968), and in turn, maintains conditions favoring this mate preference. These results suggest that the evolution of disassortative preferences is likely to depend on viability selection acting at the cue locus but also on feedbacks between cue polymorphism and mate choice. This is now calling for a mathematical framework providing general predictions on the selection regimes enabling the emergence of disassortative mating and highlighting the feedback of sexual selection on the evolution of disassortative mating when this behavior is common.

We therefore analytically explore the conditions enabling the evolution of disassortative mating by adapting a previous model of evolution of assortative mating developed by Otto et al. (2008). The model assumes a population of diploid individuals with two key loci: the first locus *C* controls variation in a single mating cue, that may be subject to viability selection. The second locus *P* controls mate preference based on the cues encoded by locus *C*. We take into account fixed and relative costs associated with choosiness. Contrary to the original model built to understand the evolution of assortative mating, alleles at preference locus *P* generate disassortative preference. Moreover, we introduce coefficients that describe the dominance at both loci to identify how the dominance relationships impact the evolution of disassortative mating.

We first analyze the model under a Quasi-linkage Equilibrium (QLE) to derive analytic expressions of changes in genetic frequency at both the cue and preference loci, providing general expectations on the conditions enabling the emergence and persistence of disassortative mating. We then use numerical simulations to explore the evolution of disassortative preferences under strong overdominant selection acting at the cue locus, that does not match the QLE assumptions. We finally compare our theoretical predictions with the few documented cases of disassortative mating and discuss why the evolution of disassortative mating may be limited in natural populations.

## Methods

Following the theoretical framework developed by Otto et al. (2008), we investigate the evolution of disassortative mating by assuming a diploid sexual species with balanced sex ratio, and considering two loci *C* and *P*. The locus *C* controls for a trait used as a mating cue and the locus *P* for the mate preference. We consider two different alleles, *a* and *b*, at locus C so that 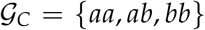 is the set of possible genotypes at this locus. This locus *C* can be under different viability selection regimes. At the mating preference locus *P*, we assume two alleles: a resident allele *M* and a mutant allele m. The set of possible genotypes at locus *P* is thus 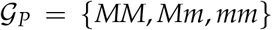. The two loci recombine with probability *r* at each birth event. We consider a discrete time model and follow the genotypes frequencies over time.

### Mating cue locus under viability selection

Dominance between the cue alleles *a* and *b* is controlled by the *dominance coefficient* at locus *C*, *h_a_*. This coefficient describes the dominance of the focal allele *a* : if *h_a_* = 0 alleles *a* and *b* are codominant and if *h_a_* = 1 (resp. −1) the focal allele *a* is dominant (resp. recessive) to *b*. If 0 < *h_a_* < 1 (resp. - 1 < *h_a_* < 0) allele *a* is incompletely dominant (resp. recessive) to *b*.

The cue induced by the genotype at locus *C* determines mating success but can also be under viability selection. We explore the evolution of disassortative mating under different viability selective regimes acting on the mating cues, specifically focusing on balancing selection regimes promoting polymorphism at locus *C*.

Let *f*(*i, k*) be the frequency of genotype 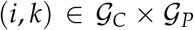. We introduce a selection coefficient *S_i_*(*f, h_a_*) acting on genotype 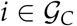, which may vary depending on genotypic frequencies at locus *C* and dominance between alleles *a* and *b*. This allows exploring different regimes of balancing selection, including negative frequency-dependent selection, that can favor polymorphism at locus *C*. Let *w_i_* be the fitness of genotype *i* resulting from viability selection acting at locus *C*

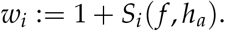

We assume that viability selection generating changes in genotype frequencies at locus *C* acts before reproduction. As a consequence, the changes in frequencies due to sexual selection depend on the frequencies at locus *C* after viability selection, described below. For 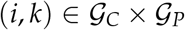:

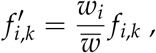

with

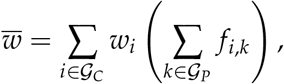

being the average fitness of the females.

### Mate choice and reproduction

Reproduction depends on the mating cues controlled by locus *C*, but also on mate preferences controlled by locus P. Each genotype 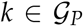 is associated with a coefficient *ρ_k_*, which quantifies how much a female of genotype *k* tends to reject males with the same cue as her own (*i.e*. the strength of disassortative preference of females). The values of *ρ_MM_* and *ρ_mm_* are fixed. For the genotype *Mm*, we introduce a dominance coefficient *h_m_* at locus *P*. Similarly to the dominance at locus *C*, this coefficient *h_m_* in [-1,1] describes the dominance of the mutant allele *m*, with the following rule:

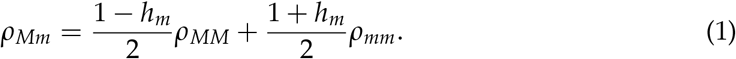

We assume females to be the choosy sex (de Cara et al., 2008; Gavrilets and Boake, 1998; Kopp and Hermisson, 2008; Lande, 1981; Otto et al., 2008), so that males can mate with any accepting females. We assume a balanced sex-ratio and consider that the frequencies of females and males with genotype *i* are equal (de Cara et al., 2008; Gavrilets and Boake, 1998; Otto et al., 2008).

To quantify the mating probability between two individuals we introduce the preference matrix *Pref*(*ρ_k_*), 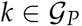, defined by:

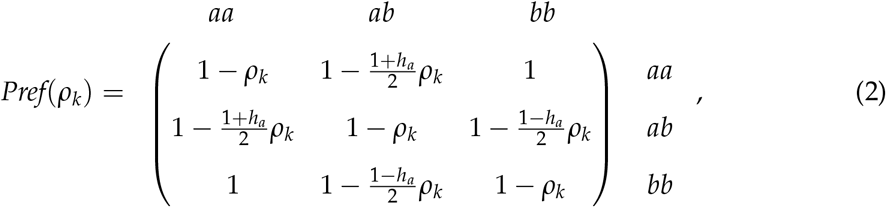

where for 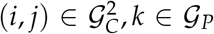, *Pref_ij_*(*ρ_k_*) measures the strength of preference of female *i* with genotype *k* at locus *P* for male *j*. With the help of this preference matrix describing disassortative mating behavior in the framework of Otto et al. (2008) (initially designed to explore the evolution of assortative mating), we investigate the evolution of disassortative mating.

For 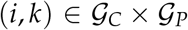, we define *T_i,k_* as the probability that a female of genotype (*i, k*) accepts a male during a mating encounter:

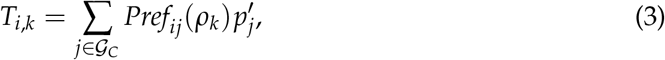

with

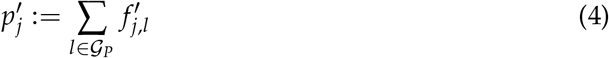

being the proportion of genotype *j* at the cue locus *C* in the population after the viability selection step.

Choosy females of genotype *k* at locus *P* are assumed to pay a fixed cost *c_f_ρ_k_* for their choosiness (the choosier a female is, the higher is this cost), that accounts for a greater investment in the search or rejection of mates. Mating behavior is indeed thought to be more costly for choosy females than for females mating with the first male encountered, regardless of displayed cue. Choosy females also pay a relative cost of choosiness, depending on the proportion of preferred males and on a coefficient *c_r_* ∈ [0,1]. This relative cost is small if the preferred mates are abundant in the population. When a female rejects a given male because he displays an unpreferred cue, she can still accept another mate with probability 1 − *c_r_*.

We define the fertility of a female of genotype 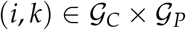 as

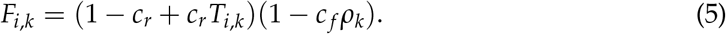

The average fertility in the population is thus:

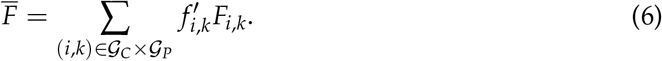

Then changes in genotypes frequencies after reproduction are as follows. For 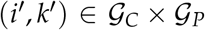:

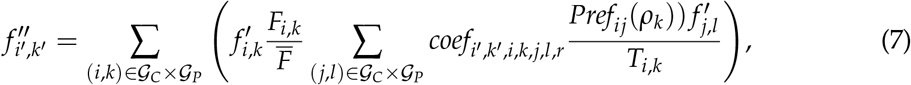

where *coef* controls the Mendelian segregation of alleles during reproduction between the choosing individual of genotype *i* at locus *C* and *k* at locus *P* and a chosen individual of genotype *j* at locus *C* and *l* at locus *P*, determining his displayed cue. The Mendelian segregation also depends on the recombination probability *r* between the cue locus *C* and the preference locus *P*. All variables and parameters used in the model are summed up in Table 1.

**Table 1:** Description of variables and parameters used in the model.

| Abbreviation | Description |
| --- | --- |
| $\mathcal{G}_C$ | Set of possible genotypes at locus C: $\mathcal{G}_C = \{aa, ab, bb\}$ |
| $\mathcal{G}_P$ | Set of possible genotypes at locus P: $\mathcal{G}_P = \{MM, Mm, mm\}$ |
| $f_{i,k}$ | Frequency of genotype $(i, k)$ in the population, $(i, k) \in \mathcal{G}_C \times \mathcal{G}_P$ |
| $f'_{i,k}$ | Frequency of genotype $(i, k)$ in the population after viability selection, $(i, k) \in \mathcal{G}_C \times \mathcal{G}_P$ |
| $f''_{i,k}$ | Frequency of genotype $(i, k)$ in the population after reproduction, $(i, k) \in \mathcal{G}_C \times \mathcal{G}_P$ |
| $p_\alpha$ | Proportion of allele $\alpha$ at locus C in the population (with $\alpha \in \{a, b\}$). |
| $p_m$ | Proportion of allele m at locus P in the population (with $m \in \{m, M\}$). |
| $r$ | Recombination probability between the loci C and P. |
| $\rho_k$ | Strength of disassortative mating within a female of genotype $k \in \mathcal{G}_P$ at locus P as described in the preference matrix (2). |
| $h_a / h_m$ | Dominance coefficient at locus C describing the dominance of allele $a / m$. |
| $S_i(f, h_a)$ | Viability selection coefficient when the allele frequencies are $f$ and the dominance coefficient at locus C is $h_a$. |
| $c_f / c_r$ | Fixed/relative cost of choosiness. |
| $D_C$ | Genetic diversity at locus C, $D_C = p_a p_b$. |
| $D_P$ | Genetic diversity at locus P, $D_P = p_m p_M$. |
| $p_{he}^{HW}$ | Proportion of heterozygotes at the Hardy-Weinberg equilibrium, $p_{he}^{HW} = 2p_a p_b$ |
| $p_{ho}^{HW}$ | Proportion of homozygotes at the Hardy-Weinberg equilibrium, $p_{ho}^{HW} = p_a^2 + p_b^2$ |
| $H_{ns} / H_{ss}$ | Heterozygote advantage due to viability/sexual selection. |
| $H$ | Heterozygote advantage. |
| $\bar{\rho}$ | Average strength of disassortative mating in the population. |
| $\Delta_\rho$ | Effect of allele $m$ on the level of disassortative mating in the population. |
| $D_{am}$ | Linkage disequilibrium between alleles $a$ and $m$ within chromosome (cis). $D_{am} = p_{am} - p_a p_m$, where $p_{am}$ is the proportion of the association between alleles $a$ and $m$ within the chromosome. |
| $D_{a,m}$ | Linkage disequilibrium between alleles $a$ and $m$ between homologous chromosomes (trans). $D_{a,m} = p_{a,m} - p_a p_m$, where $p_{a,m}$ is the proportion of the association between alleles $a$ and $m$ between homologous chromosomes. |
| $D_{he}$ | Excess of heterozygotes at locus C, $D_{he} = 1 - p_{aa}^2 - p_{bb}^2$. |
| $D_{he,m}$ | Trigenic disequilibrium measuring the association between allele $m$ and the excess of heterozygotes at locus C. |
| $\delta$ | Fitness reduction in homozygotes in numerical simulations. |
| $\mu$ | Asymmetry in viability selection acting on the two homozygous genotypes in numerical simulations. |

### Model exploration

#### QLE approximation exploring the evolution of weak disassortative preference

We use the QLE analysis results presented in a previous model of evolution of assortative mating (see Appendix B in (Otto et al., 2008)). This approach is valid when the selection coefficients, the strength of choosiness as well as costs of assortment are small; namely, for all 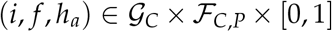 (where 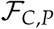 denotes the space of frequencies on 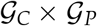) and *k* in 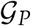, *S_i_* (*f, h_a_*), *ρ_k_*, *c_r_* and *c_f_* are of order *ϵ* with *ϵ* small. Under this hypothesis the genetic association (linkage desequilibria and departures from Hardy-Weinberg) are small (of order *ϵ*). This approach allows to obtain mathematical expressions of allele frequency changes at the cue and preference loci from the Hardy-Weinberg equilibrium. This method highlights the key evolutionary mechanisms shaping the evolution of allele frequencies at these loci. In particular, we assume that the mutant allele *m* increases disassortative preference (i.e. *ρ_mm_* > *ρ_MM_*), and investigate the evolutionary forces acting on this allele. The QLE approximation assumes a weak viability selection at the cue locus *C* and is mostly relevant to explore the evolution of weak tendency to disassortative mating (low values of *ρ*).

#### Numerical simulations

We then use numerical simulations to explore the evolutionary stable level of strength of disassortative mating when the hypothesis of weak selection is relaxed. We specifically focus on a realistic case of viability selection promoting polymorphism at the cue locus, assuming overdominance. We explore the effect of variations in key parameters, in the range where the QLE analysis is not relevant.

To explore the evolution of disassortative mating acting on the cue locus submitted to overdominance, we model a viability selection regime favoring heterozygotes. We thus set the selection coefficients associated with the different genotypes at the cue locus as:

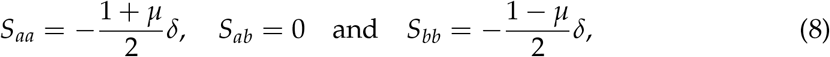

where *δ* is the fitness reduction in homozygotes and *μ* is the asymmetry in viability selection acting on the two homozygous genotypes. If *μ* = 1 (resp. −1), the disadvantage is applied to genotype *aa* (resp. bb) only, and if *μ* = 0 the disadvantage is the same for both homozygotes. To study the evolutionary stable level of strength of disassortative mating, we numerically compute the invasion gradient. First we consider a population without mutant (*p_m_* = 0), for each value of the strength of disassortative mating of the resident *ρ_MM_*, we let the initial population evolve until the genotype frequencies at the cue locus *C* reach equilibrium. At equilibrium, we introduce the mutant allele *m* with an initial 0.01 frequency. We call Δ^100^*p_m_* the change in the mutant frequency after hundred generations. We then numerically estimate

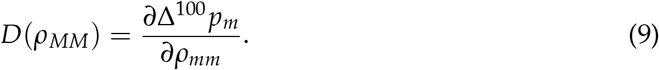

The evolutionary stable level of strength of disassortative mating is the value *ρ* for which *D*(*ρ*) = 0.

We explore the effect of variations of each of the key parameters (*δ*, *h_a_*, *μ*, *c_f_* and *c_r_*) using independent simulations. The default values for the remaining parameters follow the assumptions: codominance at cue locus *h_a_* = 0, *δ* = 1, pure symmetry in viability selection *μ* = 0 and low cost of choosiness *c_f_* = *c_r_* = 0.005. We assume no recombination *r* = 0 and codominance at preference locus *h_m_* = 0.

## Results

### Sexual selection at the cue locus generated by disassortative mating

Following the QLE approach (*i.e*. assuming that terms of the form *S_i_*(*f, h_a_*), *ρ_k_*, *c_r_* and *c_f_* are of order *ϵ* and *ϵ* is small (see Section Methods)), the change in frequency of allele *a* at the locus *C* controlling mating cue is (see Eq. (B2a) in (Otto et al., 2008)):

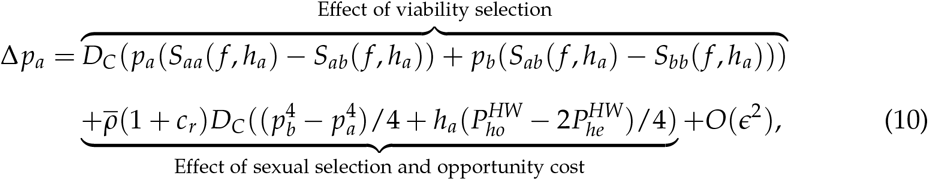

where *D_C_* = *p_a_p_b_* is the genetic diversity at locus *C*,

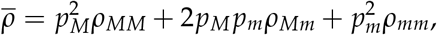

is the average disassortative mate preference at locus P,

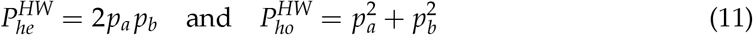

are respectively the proportion of heterozygotes and homozygotes at the Hardy-Weinberg equilibrium. Under the QLE assumption the departure from the Hardy-Weinberg equilibrium is small, hence the proportions of heterozygotes and homozygotes are close to 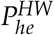 and 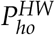.

Eq. (10) highlights that the dynamics of the mating cue allele *a* can be affected by viability and sexual selections on males and relative cost of choosiness impacting females. Contrary to assortative mating that generates positive frequency-dependent sexual selection, disassortative preferences generate negative frequency-dependent sexual selection on cue alleles (see arrows C and E in Fig. 1). The strength of this sexual selection then depends on the average strength of disassortative preference 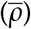. Disassortative mating also generates a relative cost of choosiness on females (see arrow D in Fig. 1). Similarly to sexual selection, this cost especially disfavors females displaying a common phenotype because these females tend to prefer males with rare phenotype.

**Figure 1:**
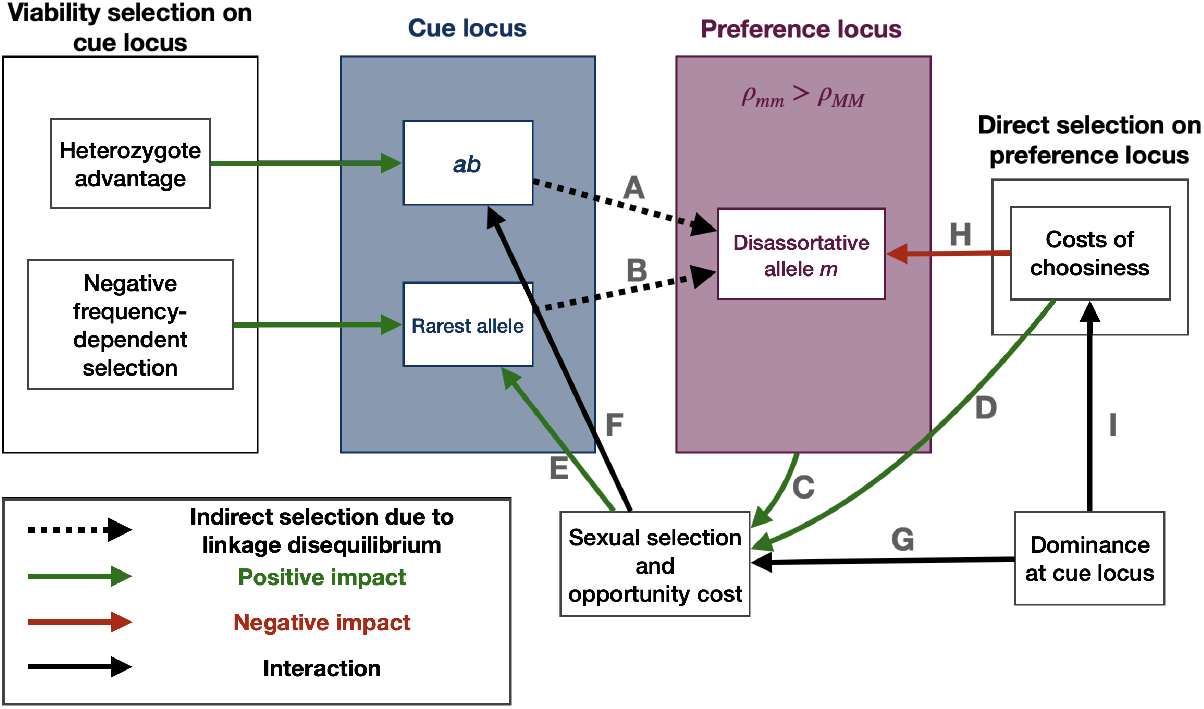
Selective forces acting on cue and preference loci. Dashed arrows represent indirect selection due to positive linkage disequilibrium between cue genotype and preference genotype. Green and red arrows represent the positive and negative impact respectively. Black arrows represent an impact that is either positive or negative (see manuscript for details). Disassortative allele is promoted by heterozygote advantage (A) and negative frequency-dependent viability selection (B) at the cue locus via indirect selection due to linkage disequilibrium. Disassortative mating triggers sexual selection on males (C) and opportunity costs on females due to a cost of choosiness (D) that generates negative frequency dependent sexual selection (E) and impacts the fitness of heterozygotes at the cue locus (F). Sexual selection often causes a disadvantage to heterozygotes at the cue locus hampering the fixation of disassortative mating. However the dominance relationship at cue locus impacts sexual selection (G). Under certain conditions sexual selection favors heterozygotes at the cue locus (C), promoting high levels of disassortative mating. The disassortative allele suffers from costs of choosiness (H). These costs depend on the dominance relationship at the cue locus (I).

Sexual selection and relative cost of choosiness also tightly depend on dominance at the cue locus *C*. When *h_a_* ≠ 0 (departure from co-dominance), the evolutionary fate of alleles is strongly influenced by their dominance. When heterozygotes are frequent at locus C, *i.e*. when allele *a* is neither rare or common (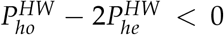 or *p_a_* ∈ (0.21,0.79), see details in Appendix A), allele *a* is favored when recessive (*h_a_* < 0), because *aa* homozygotes then display the rarest phenotype and therefore benefit from an improved reproductive success. By contrast, when heterozygotes are rare at locus *C* 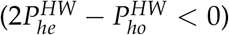, allele *a* is favored when dominant (*h_a_* > 0). Indeed, when allele *a* is rare (*p_a_* < 0.21,), *bb* individuals are numerous and preferentially mate with individuals displaying the phenotype encoded by allele *a* (the rare phenotype). Therefore, when *a* is dominant, *ab* individuals benefit from a greater mating success than *bb* individuals, thereby increasing the frequency of allele a. When the cue allele *a* is common (*p_a_* > 0.79), the dominance of allele *a* limits the reproductive success of the few remaining heterozygotes *ab* displaying the frequent phenotype shared with homozygotes *aa*, which leads to the gradual elimination of the alternative allele *b*.

These conclusions are drawn from the QLE approximation, and are relevant for moderate levels of disassortative mating (low values of *ρ*). Stronger levels of disassortative mating may lead to contrasted outcomes, because some crosses (e.g. *aa × aa*) will occur at very low frequency.

### Evolutionary fate of disassortative mating mutants

To understand the conditions enabling the evolution of disassortative mating, we now approximate the change in frequency of the mutant allele *m* at the preference locus *P*, associated with an increased level of disassortative preference as compared to the resident allele *M*. The QLE analysis highlights that the evolution of disassortative mating depends on (1) the heterozygote advantage, (2) the genetic variation at the cue locus C, and (3) the costs of choosiness, described by the terms Δ*^he^p_m_*, Δ*^C^p_m_* and Δ*^cost^p_m_* respectively. Assuming that *ϵ* is small, we get (see Eq. (B3a) in (Otto et al., 2008)):

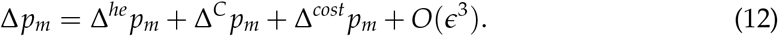

In the following sections we define these three terms and dissect the evolutionary mechanisms acting on preference alleles.

#### Disassortative mating is promoted by heterozygote advantage at the cue locus

The impact of heterozygote advantage on the frequency of the mate choice allele *m* is given by:

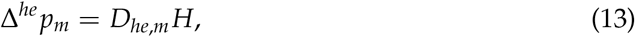

where *D_he,m_* (see (15)) is the trigenic disequilibrium describing the association between the mutant *m* at the mate choice locus *P* and heterozygotes at the cue locus *C* and *H* is the heterozygote advantage at the cue locus *C* (see (14)). The fitness advantage of heterozygotes *H* can be influenced by both viability and sexual selections, as detailed below:

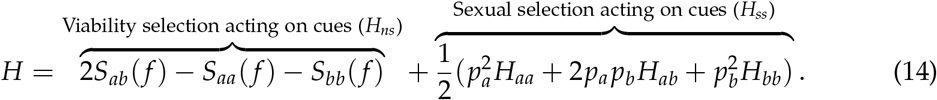

The sexual selection promoting heterozygotes at the cue locus *C* depends on mate preferences for heterozygotes over homozygotes expressed by the different genotypes 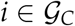 at locus *C* (*H*_i_):

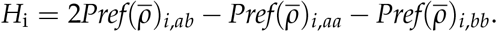

The effect of heterozygote advantage at the cue locus *C* on the disassortative mating allele *m* is then modulated by the association between the mutant *m* and heterozygotes at the cue locus (i.e. the trigenic disequilibrium *D_he,m_*), as described by Eq. (13). At QLE, the trigenic disequilibrium satisfies:

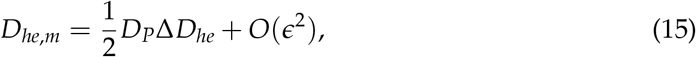

where *D_he_* is the excess of heterozygotes at locus *C* due to allele *m* and *D_P_* = *p_M_ p_m_* is the genetic diversity at locus *P*.

The trigenic disequilibrium depends on the change in the excess of heterozygotes due to allele *m* following a single round of mating. This change depends on (1) the fraction of homozygotes at the cue locus *C*, determined by allele frequencies (*p_a_* and *p_b_*) and dominance relationships (*h_a_*) and (2) the increase in disassortative preferences in the population Δ_*ρ*_ (Eq. (16)).

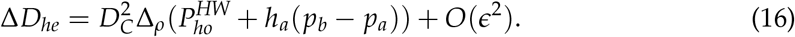

The increase in disassortative preferences Δ_*ρ*_ depends on the effect of the mutant *m* at the preference locus *P* and its frequency (Eq. (17)).

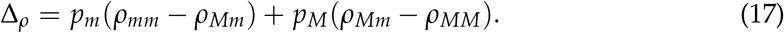

The change Δ*D_he_* has the same sign than the increase in disassortative preferences Δ_*ρ*_ (see Appendix A for details). As the mutant *m* increases the strength of disassortative preferences (*i.e. ρ_mm_* > *ρ_MM_*), Δ*D_he_*. > 0, meaning that individuals with disassortative preferences tend to produce more heterozygotes at locus *C*. As a consequence, mutant alleles *m*, increasing disassortative preferences, are preferentially associated with heterozygotes at the cue locus *C*. The disassortative mutant *m* is thus promoted when viability and sexual selections both favor heterozygotes at the mating cue locus *C* (see arrow A in Fig. 1). This contrasts with the assortative mating model of Otto et al. (2008), where the assortative allele is preferentially associated with homozygotes at cue locus, suggesting that assortative mating can be promoted when homozygotes are favored.

Dominance relationships affect the change in the frequency of heterozygotes. For instance when a rare cue allele is dominant, a round of moderate disassortative mating (i.e. *ρ_MM_* and *ρ_mm_* are small) produces more heterozygotes than when the cue allele is recessive, because the expression of the rare mating cue in heterozygotes is promoted by disassortative mate preferences.

#### Sexual selection produced by disassortative mating generates a heterozygote disadvantage limiting the evolution of such a behavior

As described above, the disassortative alleles *m* tend to be preferentially associated with heterozygotes at locus *C*. Because *ab* heterozygotes with disassortative preferences (*i.e*. carrying a *m* allele) mate preferentially with either of the *aa* or *bb* homozygotes (depending on the dominance relationship), the evolution of disassortative preferences is likely to generate a sexual selection disfavoring heterozygotes at locus *C*. This mechanism may hamper the fixation of allele *m* and may limit the evolution of disassortative mating in natural populations. This effect is determined by the mating success of heterozygotes at locus C. From Eq. (14), this sexual selection term can be written as:

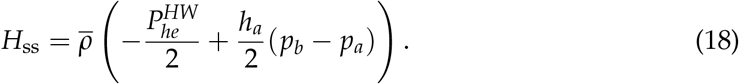

Sexual selection on heterozygotes depends on the strength of disassortative mating 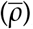, the allele frequencies at locus *C* (*p_a_* and *p_b_*) and the dominance of allele a (*h_a_*). Assuming codominance at cue locus (*h_a_* = 0), sexual selection always disfavors heterozygotes at the cue locus (see arrow F in Fig. 1). The more common disassortative preferences are in the population, the higher this sexual selection acting against heterozygotes is. Since the disassortative allele *m* is preferentially associated with heterozygotes at cue locus, it suffers from sexual selection caused by disassortative mating. The spread of a disassortative allele is thus limited by this negative feedback.

However, the sexual selection acting against heterozygotes at the cue locus depends on the dominance relationship at the cue locus (see arrow G in Fig. 1). Assuming strict dominance at the cue locus (*h_a_* = −1 or *h_a_* = 1), heterozygous individuals are indistinguishable from homozygotes, therefore modifying the proportion of phenotypes in the population. Heterozygote advantage at the cue locus due to sexual selection increases when the most common allele is recessive: when allele *a* is recessive and common heterozygous males *ab* have the same phenotype as homozygotes *bb*. *ab* males then display the rarest phenotype and benefit from negative frequency-dependent selection. When the dominant cue allele is sufficiently rare, sexual selection favors heterozygotes (see Appendix A), generating a positive feedback loop favoring the evolution of disassortative mating (see arrow F in Fig. 1). However, this effect should often be transient because negative frequency-dependent sexual selection rapidly balances phenotypic cue frequencies. In the general case where allele frequencies are balanced at the cue locus, sexual selection is thus expected to limit the evolution of disassortative mating.

Sexual selection also impacts the evolution of assortative mating (Otto et al., 2008), where the assortative allele is preferentially associated with homozygotes at the cue locus. Similarly to disassortative mating, sexual selection is often thought to limit the evolution of assortative mating. However homozygote disadvantage due to assortative mating decreases with the proportion of homozygotes in the population. Assortative mating promotes homozygotes, this preference may thus suffer from a weak negative feedback loop, in contrast with the evolution of disassortative mating.

#### Disassortative preferences are favored when the rarer allele is promoted

The change in the frequency of cue alleles impacts the evolution of preference alleles. This impact is described by the term:

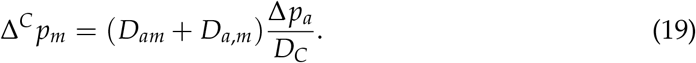

As highlighted in Eq. (19), the invasion of a disassortative mutant *m* depends on its linkage with the cue allele *a* (either in *cis* or in *trans*, described by *D_am_* and *D_a,m_* respectively) and on the variation in the frequency of allele *a* (Δ*p_a_*). If allele *m* is associated with allele *a*, the frequency of allele *m* increases with the rise of frequency of allele *a*. The QLE approximates the *cis* and *trans* linkage desequilibria between the mutant allele *m* and the cue allele *a* as:

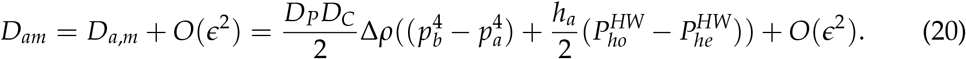

*D_am_* and *D_a,m_* have the same sign as *p_b_* − *p_a_* (see Appendix A for more details), thus *D_am_* and *D_a,m_* are positive (resp. negative), when allele *a* is the rarer (resp. most common). Contrary to assortative alleles preferentially associated with the most common cue allele (Otto et al., 2008), Eq. (20) indicates that the disassortative mating allele *m* tends to be linked with the rarer allele at locus *C*. This predicts that disassortative mating is likely to emerge when viability selection on the cue provides fitness benefit to rare alleles (see arrow D in Fig. 1), while assortative mating is promoted when the most common cue alleles are favored.

Disassortative allele *m* also tends to be more tightly linked either to the dominant cue allele when the frequency of homozygotes is high, or to the recessive allele when the frequency of heterozygotes is high (*i.e* when 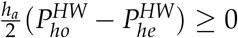), increasing the association between alleles *a* and *m*). The effect of dominance can thus modulate the association between allele *m* and the rarer cue allele.

Given that (1) the disassortative allele *m* is associated with the rarer cue allele and (2) disassortative mating promotes the rarer allele via sexual selection, the disassortative mating allele *m* could benefit from a positive feedback loop promoting the evolution of disassortative mating. However, negative frequency-dependent sexual selection rapidly increases the frequency of the initially rare allele, limiting the spread of the *m* allele in the population. The initially rarer allele may become as common as the other allele breaking the linkage disequilibrium between allele *m* and alleles at cue locus. Thus this positive effect of sexual selection on the evolution of disassortative mating could be broken with the increase of the initially rarer allele frequency.

#### The costs of choosiness limit the fixation of disassortative mating

The evolution of mate preferences is generally limited by the costs associated with choosiness. Eq. (21) shows that both fixed and relative costs of choosiness indeed limit the fixation of the disassortative mutant *m* (see arrow H in Fig. 1):

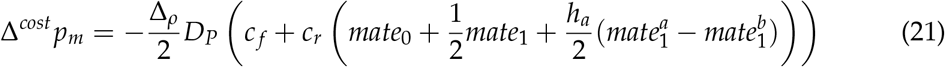

where *mate_i_*, *i* ∈ {0,1} and 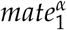, *α* ∈ {a,b} describe the proportion of mating partners sharing different numbers of alleles (see Eq. (22) and (24)). The costs of choosiness disfavor preference alleles increasing disassortative choices (i.e. when *ρ_mm_* > *ρ_MM_*) (see Appendix A for details). The relative cost of choosiness then crucially depends on the proportion of preferred mates. This effect can be captured by the parameters *matek*, *k* ∈ {0,1} representing the probability that a female encounters a male differing by *k* allele at locus *C* at the Hardy-Weinberg equilibrium:

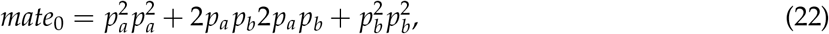

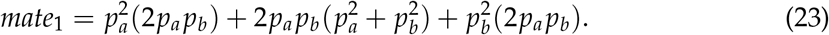

The mating between individuals differing by zero (*mate*_0_) or one cue allele (*mate*_1_) may be partially avoided when individuals have a disassortative preference, resulting in a cost *c_r_* for the choosy female that may fail to find a suitable male. The term 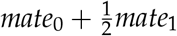 is minimal when *p_a_* = *p_b_*, so that the impact of the relative cost of choosiness is weaker when the cue alleles are in similar proportions in the population, maximizing the opportunities for females to find a male displaying the preferred cue. The dominance at the cue locus *C* then modulates the crosses at the Hardy-Weinberg equilibrium between individuals carrying at least one allele 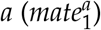 and between individuals carrying at least one allele 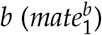

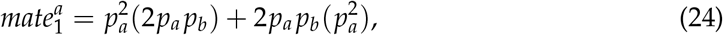

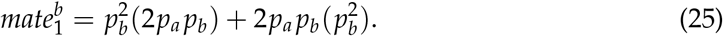

When *a* is dominant (*h_a_* > 0), matings between individuals sharing at least one allele 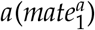 are limited by disassortative preference, leading to an increased cost of choosiness. By contrast, matings between individuals sharing at least one allele 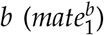 are promoted by disassortative preference, therefore limiting the cost of choosiness. The difference between 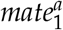 and 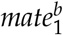 is thus crucial to understand the impact of the dominance relationship at locus *C* on the cost of choosiness. This difference is given by:

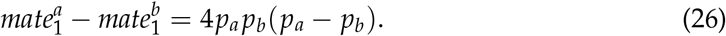

Thus when *a* is dominant (*h_a_* > 0), the relative cost of choosiness is limited when allele *a* is rare, because *bb* homozygotes will frequently meet *ab* heterozygotes displaying their preferred cue. Symmetrically, the cost of choosiness acting on the mutant allele *m* is higher when the most common cue allele is dominant. The dominance relationship therefore influences the evolution of disassortative mating also by modulating the costs of choosiness (see arrow I in Fig. 1).

#### Recombination rate does not impact the evolution of disassortative mating based on a matching rule

The QLE approximation revealed no effect of the recombination rate *r* between cue and preference alleles, suggesting that it does not impact the evolution of disassortative mating. Similarly, recombination does not impact the analytical results brought by QLE approach applied to the evolution of assortative mating (Otto et al., 2008). These two models assume mate preferences based on matching rule, *i.e*. that females use their own cue to choose their mate (Kopp et al., 2018). Under this assumption, a mutant allele *m* immediately translates into disassortative mating in any female carrying it, independently from her genotype at the cue locus. By contrast, assuming a trait/preference rule, *i.e*. when females choose their mate independently of their own cue, any preference allele in a female does not always generate a disassortative behaviour, depending on her genotype at the cue locus. Under such a preference/trait hypothesis, the recombination rate would likely impact the evolution of disassortative preference.

### Evolution of disassortative mating assuming strong overdominance at the cue locus

The QLE approximation allows to draw analytic approximations for the change in frequencies at both loci, assuming low levels of selection. Appendix B shows that QLE approximations are relevant when the parameters *S_i_*(*f, h_a_*) for all 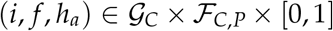, *ρ_i_* for all 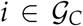, *c_r_* and *c_f_* are small, but are not valid outside these conditions. Since, we could not perform a local stability analysis using analytical derivation, we run numerical simulations to study ecological situations where viability selection at the cue locus can be strong and/or marked mate preferences lead to high rate of disassortative mating.

Well-documented cases of disassortative mating in natural population present strong heterozygote advantage (Jay et al., 2019; Tuttle et al., 2016). We thus focus on the evolution of disassortative mating acting on a cue locus where strong overdominance is operating (Fig. 2).

**Figure 2:**
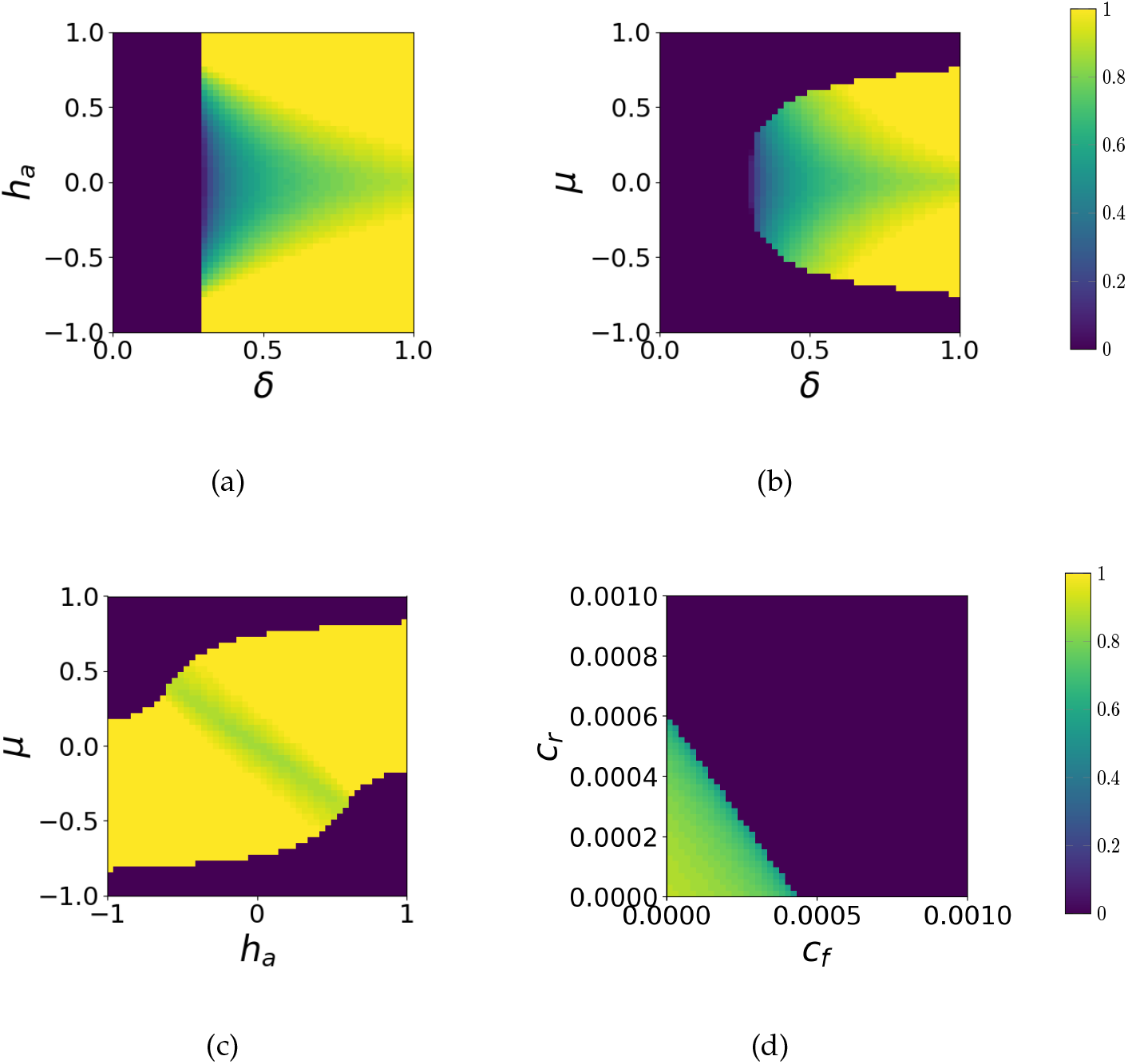
Evolutionary stable level of strength of disassortative mating *ρ* acting on a cue locus submitted to overdominance. We plotted the evolutionnary stable level of strength of disassortative mating *ρ*. The effects of key parameters on the evolution of the disassortative mating acting on the cue loci submitted to overdominance are explored in the different panels: (a) Effect of fitness reduction in homozygotes *δ* and of dominance coefficient at the cue locus *C h_a_*, (b) Effect of fitness reduction in homozygotes *δ* and asymmetry in this reduction on the two homozygotes *μ*, (c) Effect of dominance coefficient at the cue locus *C h_a_* and asymmetry in the fitness reduction on the two homozygotes *μ* and (d) Effect of fixed cost of choosiness (cf) and relative cost of choosiness (*c_r_*). The default parameters values are as follows: *h_a_* = *h_m_* = 0, *r* = 0, *δ* = 0.9, *μ* = 0 and *c_r_* = *c_f_* = 0.005.

#### Disassortative mating is favored by asymmetrical overdominance

Our simulations show that the difference between the fitness of heterozygotes and homozygotes has a strong effect on the evolution of disassortative mate preferences (Fig. 2(a) and 2(b)). Higher levels of disassortative mating are favored when heterozygotes at the cue locus are advantaged by viability selection (i.e. when homozygotes suffers from a significant genetic load *δ*, Fig. 2(a) and 2(b)), consistent with the predictions brought by the QLE approximation. Interestingly, higher levels of disassortative mating are favored when there is a moderate asymmetry (*μ*) in the negative selection acting on homozygotes at the cue locus, *i.e*. when one out of the two cue alleles is associated with a stronger genetic load (Fig. 2(b)). Selection indirectly acting on mating preference indeed crucially depends on genotypic frequencies at the cue locus *C*, which become unbalanced under asymmetrical selection. Unbalanced cue allele frequencies tend to increase the frequency of homozygotes compared to the frequency of heterozygotes, increasing the relative advantage of heterozygotes due to viability selection, to sexual selection and to opportunity cost. As disassortative preference tends to be linked with heterozygotes, high levels of disassortative mating are favored by the unbalanced cue allele frequencies.

Because disassortative mating mutants are preferentially associated with the rare allele (carrying the recessive genetic load), once the asymmetrical selection against the rare allele is too strong, it prevents the emergence of the disassortative mating alleles associated with this maladaptive cue allele. When the negative viability selection on the rare allele is lower than a threshold, viability selection allows the emergence of the disassortative mating mutant and even favors the evolution of stronger levels of disassortative mating because as the level of disassortative behavior increases, the disadvantage of being associated with the rarer allele becomes weaker.

Asymmetrical overdominance therefore promotes the evolution of disassortative mating preference, but only when the asymmetry in the genetic load associated with cue alleles is not too high.

#### Interactions between dominance and fitness of cue alleles determine the evolution of disassortative mate preferences

High levels of disassortative mating are favored when dominance relationships at the cue locus are strict (i.e. when allele *a* (resp. *b*) is fully dominant to *b* (*h_a_* = 1) (resp. *a* (*h_a_* = −1)) as highlighted on Fig. 2(a). The dominant allele is disfavored by sexual selection generated by disassortative mating. When the dominant allele is rare the association of disassortative preference and cue heterozygosity increases, promoting high levels of disassortative mating. Moreover when the dominant allele is rare, the impact of the costs of choosiness on frequency changes is lower, further promoting high levels of disassortative mating.

When combining both effects leading to unbalanced cue allele frequencies (i.e. dominance and asymmetrical negative selection on cue alleles), we show that high levels of disassortative mating are strongly favored when the fitness reduction in homozygotes is associated with the dominant cue allele (Fig. 2(c)). This numerical result is consistent with the prediction drawn from the QLE approximation, because in this case, the dominant allele is in low frequency (because of both viability and sexual selections).

#### The challenging evolution of disassortative mating

Numerical simulations confirm that the evolution of disassortative mating is challenging when moderate overdominance (enhancing the fitness of heterozygotes) is at play at the cue locus. In most cases, strict disassortative mating is not favored. The higher the disassortative preferences, the more sexual selection acts against heterozygotes. When heterozygote advantage is not strong enough, sexual selection caused by mating preferences can overcome heterozygote advantage, favoring intermediate level of disassortative mating (see green areas on Fig. 2(a) and 2(b)). By contrast, when viability selection produces strong heterozygote advantage (*δ* is high) that can compensate sexual selection, complete disassortative preferences can be fixed (see Fig. 2(a) and 2(b)).

The costs of choosiness may further limit the evolution of the disassortative mutant. Fig. 2(d) shows that disassortative mating is under positive selection only when the costs of choosiness are limited (at least inferior to 0.03).

## Discussion

### Predicted selection regimes promoting disassortative mating match empirical observations

Our results show that disassortative mating is promoted either (1) when heterozygotes at cue locus are in average fitter that homozygotes or (2) when viability selection on cue favors the rarest cue allele. These selection regimes promoting disassortative mating are opposed to the selection regimes promoting assortative mating, such as homozygote advantage at cue locus or viability selection on cue favoring the most common allele (Otto et al., 2008) (see Table 2).

**Table 2:** Comparison between the evolution of disassortative mating based on the present study and the evolution of assortative mating based on Otto et al. (2008)’s study.

|  | Present model studying the evolution of disassortative mating | Otto et al. (2008) model studying the evolution of assortative mating |
| --- | --- | --- |
| Viability selection on mating cue that promotes preferences | Heterozygotes advantage<br>Negative frequency-dependent viability selection | Homozygotes advantage<br>Positive frequency-dependent viability selection |
| Sexual selection on mating cue due to preferences | Is expected to disadvantage heterozygotes unless when one type of homozygote is common and heterozygotes display the same mating cue as rare homozygotes<br>Negative frequency-dependent sexual selection | Is expected to disadvantage homozygotes unless when females sufficiently reject males differing by one cue allele and homozygotes are common<br>Positive frequency-dependent sexual selection |
| Relative cost of choosiness | Lower when one type of homozygote is common and heterozygotes display the same mating cue as rare homozygotes | Lower when one type of homozygote is common |

Interestingly, our simulations also show that higher levels of disassortative mating are promoted when one cue allele is dominant. The dominance relationship can indeed decrease sexual selection and relative cost of choosiness impairing the evolution of disassortative preferences.

Simulations also highlight that higher levels of disassortative mating are promoted when the dominant allele is disfavored when homozygous. This effect is consistent with the observed cases of disassortative mating. For instance the butterfly *H. numata* displays a strong disassortative mating based on wing-pattern phenotype (in a tetrad experiment, 3/4 of the realized crosses were involving disassortative pairs) (Chouteau et al., 2017). In this species, the variation in wing-pattern morphs is controlled by a supergene with three main haplotypes (Joron et al., 2011). The dominant haplotypes are associated with a low survival of homozygous larvae (Jay et al., 2019). This case of disassortative mating seems to gather the conditions pinpointed by our model to enable the evolution of higher levels of disassortative mating.

Similarly, in the white-throated sparrow *Zonotrichia albicollis* an almost strict disassortative mating based on plumage morphs (*white* or *tan*) has been reported (Throneycroft, 1975). Two supergene haplotypes, here refered to as *t* and *w*, control this variation in plumage coloration. Individuals with *tt* genotype have a *tan* coloration whereas individuals carrying *tw* and *ww* genotypes have a *white* coloration. However the dominant haplotype *w* is associated with strong genetic load, generating homozygote disadvantage in *ww* individuals (Tuttle et al., 2016). Individuals with *white* coloration may be advantaged over *tan* individuals because they invest less into parental care (Knapton and Falls, 1983), generating an advantage of heterozygotes *tw* over homozygotes tt. Here the dominant cue allele is again associated with a strong disadvantage when homozygous, which, according to our results, strongly favors the emergence of disassortative preferences (see Fig 3).

**Figure 3:**
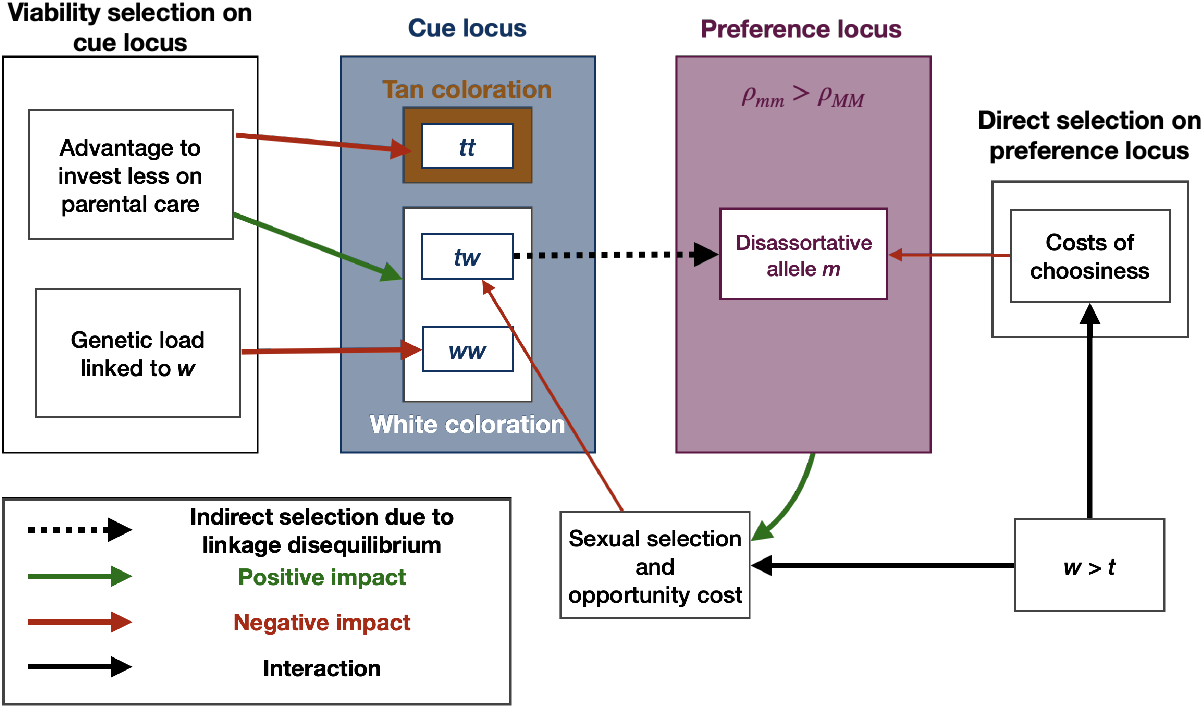
Selective forces acting on cue loci in the example of the white-throated sparrow (*Zonotrichia albicollis*). See Figure 1 for details of the meaning of symbols.

### Polymorphism at the mating cue has a crucial effet on the evolution of disassortative mating

The number of mating cues within the population is an important parameter in the evolution of mate preference (Otto et al., 2008), because it modulates the opportunity costs generated by choosiness. In our model, we consider only two cue alleles, generating at most three different cue phenotypes in the population (phenotypes displayed by individuals *aa, ab* and bb). With a higher number of alleles, the number of phenotypes would be greater. Under disassortative mating, these phenotypes should have their frequencies balanced by negative frequency-dependent selection. Thus both females and males would still have sufficient mating opportunities, weakening the relative cost of choosiness and sexual selection. Then disassortative mating should evolve more easily when the number of mating cue is higher. This may have favored the evolution of disassortative preference targeting MHC loci, where multiple alleles are maintained by selection (de Vries, 1989).

When the mating cue is a quantitative trait (e.g. size-related preferences, (Janicke et al., 2019; Jiang et al., 2013)), variations within populations may be considered as multiple cues, depending on the discrimination rules of the choosy partners. If quantitative variations are perceived as multiple differentiated phenotypes, it would probably promote the evolution of disassortative mating, in a similar manner as high level of discrete polymorphism.

The number of mating cues maintained within a population can also be increased via contacts between populations. The effect of immigration of individuals displaying alternative cues on the evolution of disassortative mating will then depend on viability selection. Cotto and Servedio (2017), show that the contact between populations promotes higher level of assortative mating, because individuals adapted to different habitats produces intermediate offspring maladaptive in each habitat. Contacts between locally adapted populations may thus limit the evolution of disassortative mating because it generates viability selection against hybrids, disfavoring such preferences.

Mating opportunities also depend on the distribution of cues in the population. A more balanced cue distribution within population often increases the negative effect of sexual selection on the evolution of assortative preferences (Otto et al., 2008). For instance, migration between populations has been shown to limit the evolution of further assortative mating because it promotes a more balanced polymorphism within populations and therefore increases the negative effect of sexual selection (Servedio, 2011). Similarly, migration between populations may limit the evolution of disassortative mating, because the resulting more balanced polymorphism increases the negative sexual selection.

### Negative feedback in the evolution of disassortative mating contrasts with the evolution of assortative mating

A striking result from our analyses stems from the role of sexual selection generated by disassortative preferences on its evolution, which contrasts with the evolutionary dynamics of assortative mating. Our results confirm that the sexual selection generated by disassortative mating often limits its own spread, as already mentioned by Nuismer et al. (2008). Indeed, the disassortative mating allele is generally associated with heterozygotes at the cue locus. Individuals with such allelic combinations tend to preferentially mate with homozygotes, generating sexual selection disfavoring heterozygotes at the cue locus. However, this sexual selection acting against heterozygotes depends on the distribution of cue allele frequency (see more details in Tab. 2).

Similarly, the evolution of assortative mating is thought to be limited by sexual selection (Otto et al., 2008) (but sexual selection can promote the evolution of assortative mating in some cases, see more details in Tab. 2). However, this negative effect of sexual selection decreases when the proportion of homozygotes at the cue locus is high. Assortative mating usually produces more homozygotes than random mating: a decrease in the level of heterozygosity at the cue locus is thus expected when assortative preferences are spreading within a population. During the evolution of assortative mating, the negative effect of sexual selection on the evolution of assortative mating decreases as the proportion of homozygote increases. The evolution of disassortative mating may therefore be more severely impaired by sexual selection than the evolution of assortative mating.

Hence, favorable conditions for disassortative preferences may result in intermediate values of choosiness in the population. In two meta-analyses (Janicke et al. (2019); Jiang et al. (2013)) covering 1,116 and 1,447 measures of strength of assortment respectively, most of the values corresponding to disassortative mating range from −0.5 to 0 (but see below exception), suggesting that intermediate values of strength of disassortative mating are frequently observed. By contrast, most values corresponding to assortative mating behavior range from 0 to 1, suggesting that the evolution of strict assortative mating is observed in a wide range of organisms.

### Alternative genetic architectures of mate preferences may limit the evolution of disassortative mating

The genetic architecture of preference may also have an impact on the evolution of disassortative mating. Theoretical studies on the evolution of assortative mating usually rely on two main types of matching rules Kopp et al. (2018): (1) when mate choice of an individual depends on its own phenotype (*matching rule*) and (2) when preference is independent from the phenotype of the chooser (*preference/trait* rule). The evolution of assortative mating is strongly promoted either when assuming the *matching rule*, or when the cue and *preference/trait* loci are tightly linked (Kopp et al., 2018). Here, our results on the evolution of disassortative mating are obtained assuming a *matching rule*, and we expect that assuming a *preference/trait rule* might limit such an evolution, because selection might break the unmatching allelic combinations. In the specific case of polymorphic mimicry, Maisonneuve et al. (2019) showed that under *preference/trait* rule, disassortative mating can emerge only if the preference and the cue loci are fully linked.

Moreover, here we only consider a single choosy sex. However, when both sexes are choosy (Servedio and Lande, 2006), the positive selection on the evolution of mate preference in one sex may be relaxed when strong mate preferences are fixed in the other sex (Aubier et al., 2019). Drift then leads to periodic cycles where male and female alternatively become the most choosy sex (Aubier et al., 2019).

### Conclusions

Our analytical and numerical results provide a general theoretical framework establishing the conditions enabling the evolution of disassortative mating. Our results pinpoint two selective regimes on mating cue that promote disassortative mating through indirect selection : heterozygote advantage and negative frequency-dependent selection. We also observe that disassortative mating generates sexual selection that often hamper its own fixation, leading to intermediate level of disassortative mating. This sexual selection depends on the dominance at the cue locus: if one type of homozygote at the cue locus is common and if heterozygotes display the same cue as the rare homozygote, sexual selection promotes the evolution of disassortative mating. We also show that this condition reduces the costs associated with choosiness. Interestingly, the favorable selective conditions predicted by our model match with two well-characterized cases of strong disassortative mating.

## Appendix A: Details of analytic results

We study the evolution of alleles frequencies in a two locus diploid model. One locus controls the mating cue (with two alleles *a* and *b*) and the other controls mate preference (with two alleles *M* and m). The model is described in the main file. We used a QLE analysis to approximate the changes in frequency of the cue allele *a* and of the preference allele *m* (see main file for more details). Here, we detail how our analytic results pinpoint some mechanisms explained in the main file.

### The dominance relationship at cue locus impacts the action of sexual selection in a way depending on the proportion of heterozygotes

The approximation of the change in frequency of the cue allele *a* using QLE analysis is given by:

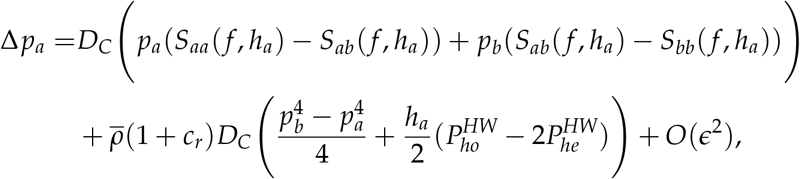

where we recall that under the QLE approximation, *ϵ* is a small quantity.

Here we aim to study the impact of the dominance relationship on the variation of allele *a* frequency. We therefore study the sign of the term

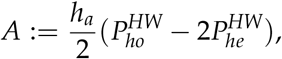

describing the effect of the dominance relationship. As 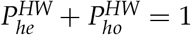 by definition we have:

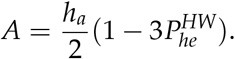

Thus *A* > 0 when *a* is partially dominant (*h_a_ >* 0) (resp. recessive (*h_a_* < 0) and the proportion of heterozygotes is lower (resp. higher) than 1/3. This entails that when the proportion of heterozygotes is low (resp. high), the dominant (resp. recessive) cue allele is favored.

The condition on 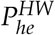 translates in a condition on *p_a_* as follows:

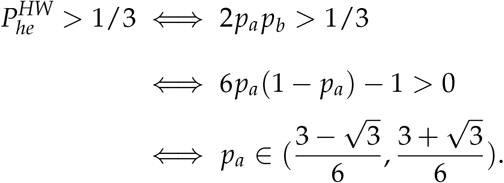

For the sake of readability we use in the manuscript the approximation 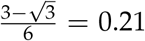 and 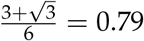.

### Disassortative preference promotes heterozygote excess at cue locus

We develop the expression of the change of excess of heterozygotes at cue locus due to the preference allele *m*, Δ*D_he_*:

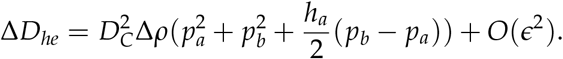

Thus Δ*Dh_e_* depends on the sign of the term 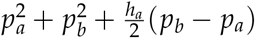. But the latter can be written as follows:

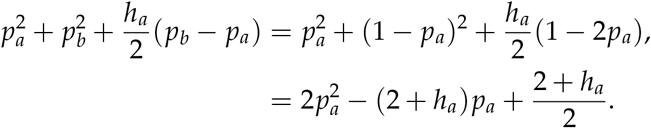

This term is a quadratic function in *p_a_*. The value of its discriminant is:

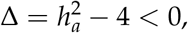

which entails that this quadratic function is always positive, and that Δ*D_he_* has the same sign as Δ*ρ.* Hence, when allele *m* is associated with higher disassortative preference (Δ*ρ*) it promotes heterozygoty excess.

### Sexual selection generated by disassortative mating can favor or disfavor heterozygotes at cue locus

The heterozygote advantage due to sexual selection is given by:

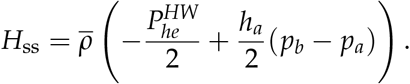

To study the impact of sexual selection on heterozygotes we look at the sign of 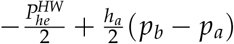. The latter can be written as follows:

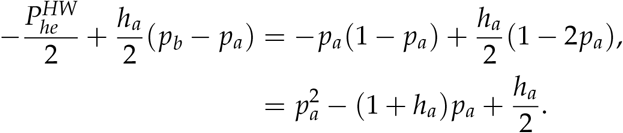

It is a quadratic function in *p_a_* with discriminant:

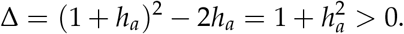

Therefore *H*_ss_ is equal to

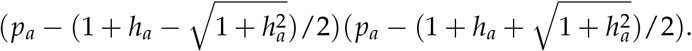

When there is codominance at cue locus (i.e. *h_a_* = 0), we have 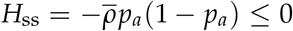, thus disassortative preference always disfavor heterozygotes at cue locus. A classical functional study yields that 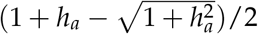 belongs to [−1,1] and has the sign of *h_a_*, and that 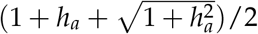 belongs to [0,1]. Asa consequence when *h_a_* ≠ 0, *H_ss_* can be either positive or negative depending on the frequency of allele *pa*. Therefore when the dominance relationship is unbalanced, sexual selection due to disassortative mating may favor or disfavor heterozygotes at cue locus.

### Mutant allele m is always associated with the rarer cue allele

The associations between allele *m* and cue alleles are given by the *cis (D_am_*) and *trans* (*D_a,m_*) linkage disequilibria. At QLE these linkages can be approximate by:

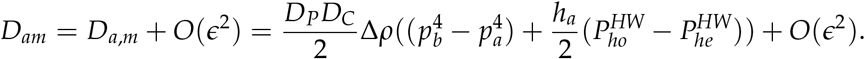

To understand the association between allele *m* and cue alleles we have to look at the sign of 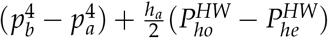. But the latter can be written as follows:

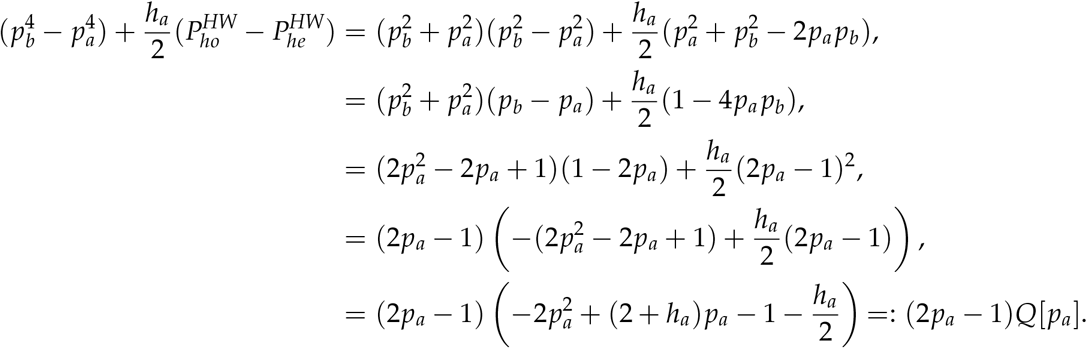

where *Q* is a quadratic function, with discriminant:

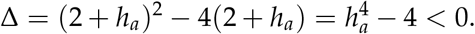

It entails that *Q* is always negative, and that *D_am_* and *D_a,m_* have the sign of 1 − 2*p_a_* (i.e. *p_b_* − *p_a_*). As a consequence preference allele *m* is associated with the rarer allele at cue locus at QLE.

### Costs of choosiness penalize the preference allele associated with higher levels of disassortative mating

Here we study the impact of the costs of choosiness on frequencies of preference alleles. We recall that the impact of the costs of choosiness on allele *m* frequency is given by:

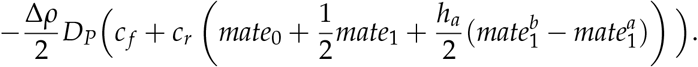

We are interested in the sign of the term *B* defined by:

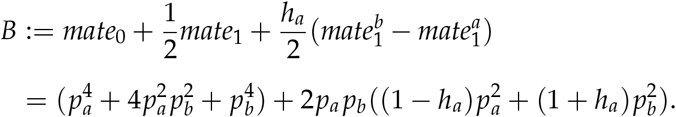

*B* is this the sum of two positive terms. Hence when Δ*ρ* is positive (i.e. when *ρ_mm_* > *ρ_MM_*), the costs of choosiness penalize the preference allele associated with higher disassortative preferences.

## Appendix B: Comparison of QLE analysis results with numerical simulations

We used a QLE analysis to draw analytic approximations for the changes in frequencies at the cue and preference loci (Δ*p_a_* and Δ*p_m_*). The results of the QLE analysis are only relevant when for all 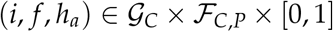 (where we recall that 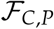 denotes the space of frequencies on 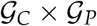), *S_i_*(*f, h_a_*) = *O*(*ϵ*); for all *k* in 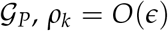, *c_r_* = *O*(*ϵ*) and *c_f_* = *O*(*ϵ*) with *ϵ* small.

To illustrate that the QLE results provide a good approximation under the QLE hypothesis, we then compare the values of the frequency changes predicted by the QLE analysis (Δ*^QLE^p_a_* and Δ*^QLE^p_m_*) with values from numerical simulations (Δ*^num^p_a_* and Δ*^num^pm*). We define 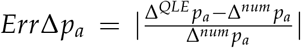 and 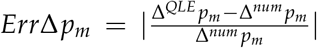 to quantify the error of the QLE approximation. We assume that the viability selection does not depend on the frequencies distribution and of the dominance at cue locus. Then for all 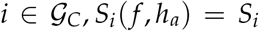, *S_i_*(*f, h_a_*) = *S_i_*. The results are plotted in Figures B1 and B2 and show that the error of the QLE approximations are low when the hypotheses of the QLE are satisfied *i.e*. the parameters *S_aa_*, *S_ab_*, *S_bb_*, *ρ_MM_*, *ρ_mM_*, *ρ_mm_*, *c_r_* and *c_f_* are small.

---

**Figure B1:**
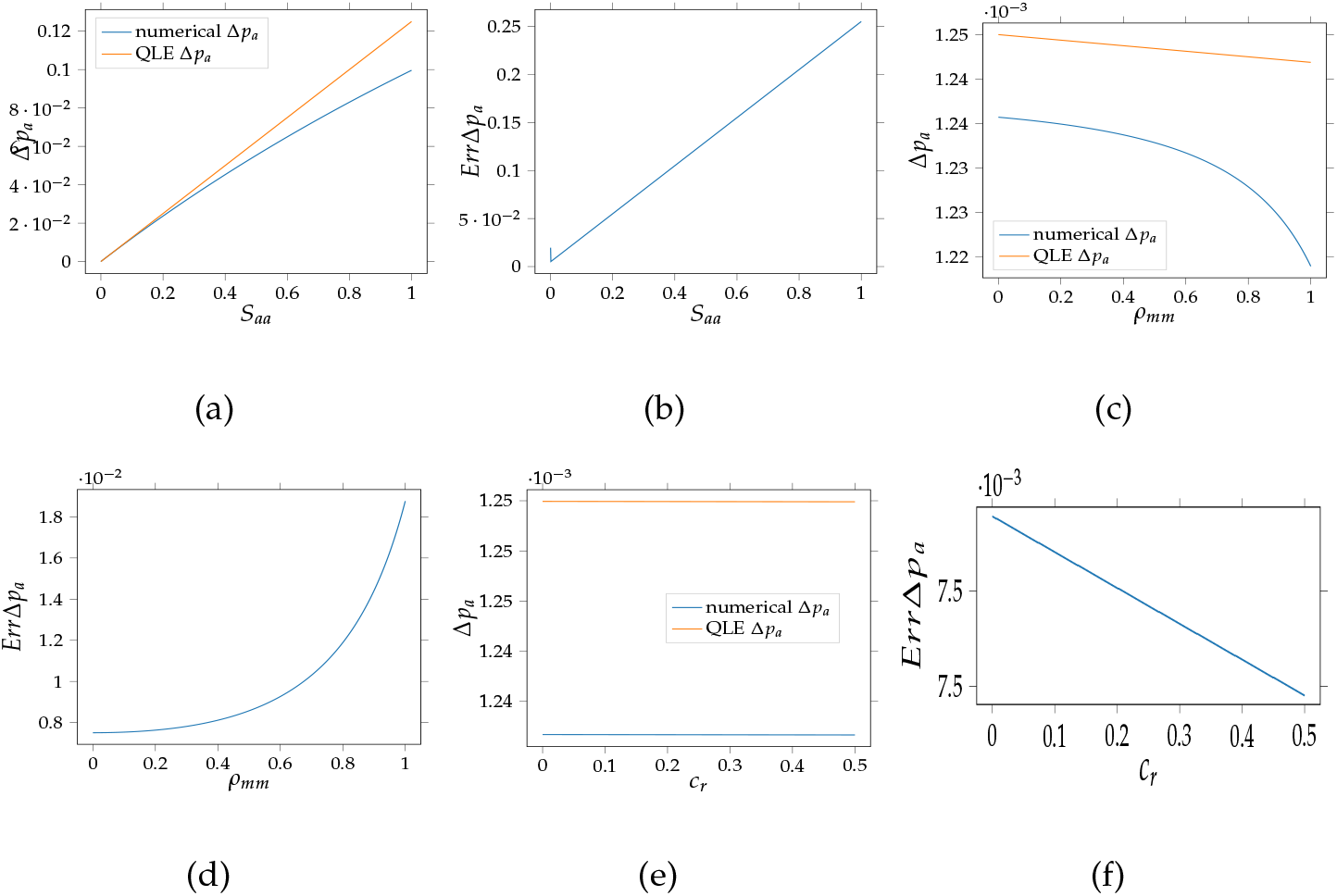
Change of allele *a* frequency at cue locus (Δ) after the introduction of mutant predicted by the QLE analysis (orange lines) and by numerical simulations (blue lines). The effect of key parameters on the evolution of the disassortative mating is explored in the different panels: (a)(b) Effect of the viability selection acting on homozygote *aa* at cue locus (*S_aa_*), (c)(d) Effect of the strength of disassortative mating within an individual of genotype *mm* at preference locus (*ρ_mm_*), (e)(f) Effect of the relative cost of choosiness (*c_r_*). The default parameters values are as follows: *ρ_MM_* = 0, *ρ_mm_* = 0.01, *h_a_* = 1, *h_m_* = −1, *r* = 0.1, *c_r_* = 0, *c_f_* = 0, *S_aa_* = 0, *S_ab_* = 0.01 and *S_bb_* = 0.

**Figure B2:**
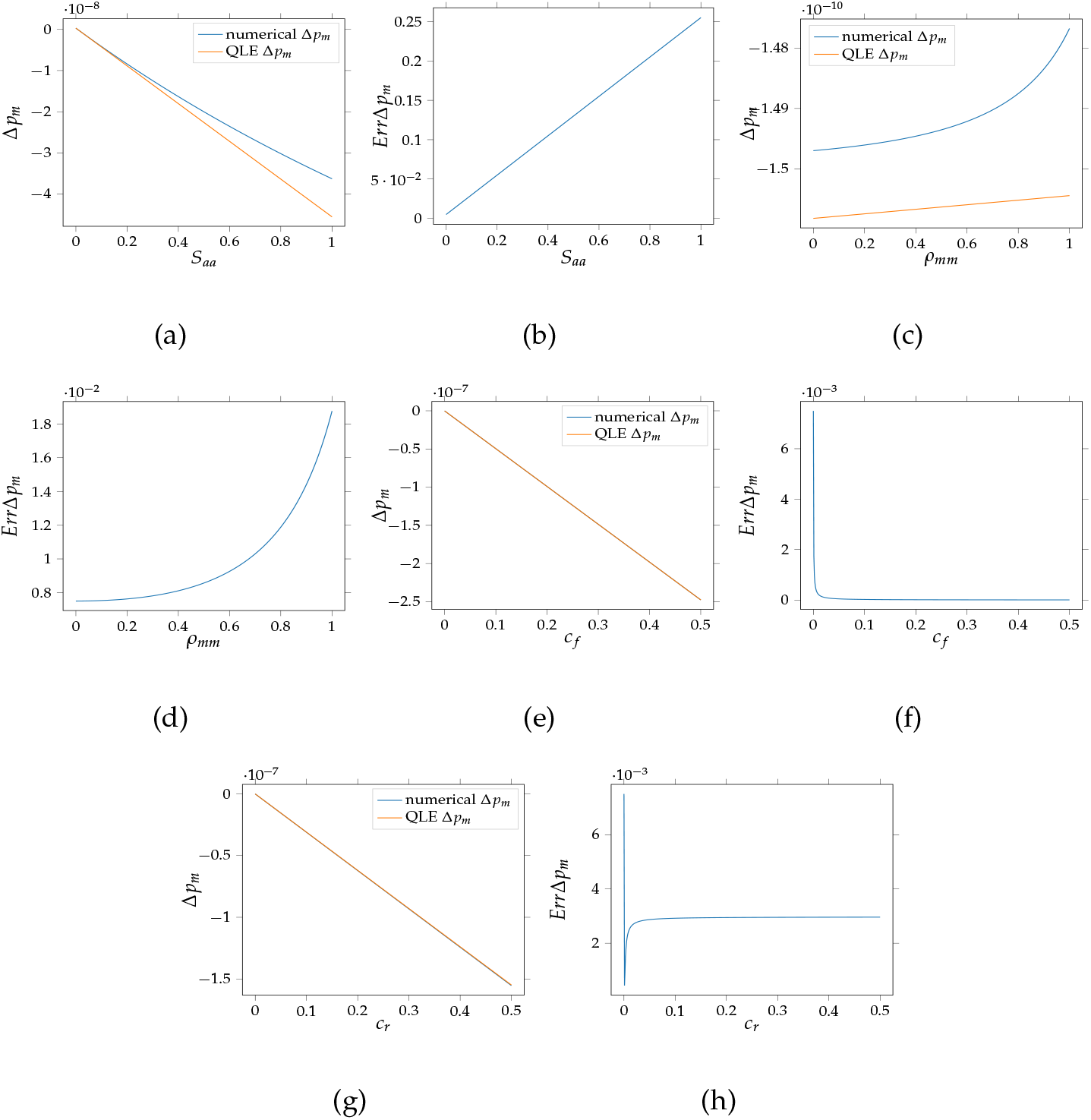
Change of allele *m* frequency at preference locus (Δ*p_m_*) after the introduction of the mutant predicted by the QLE analysis (orange lines) and by numerical simulations (blue lines). The effect of key parameters on the evolution of the disassortative mating is explored in the different panels: (a)(b) Effect of the viability selection acting on homozygote *aa* at cue locus (*S_aa_*), (c)(d) Effect of the strength of disassortative mating within an individual of genotype mm at preference locus (*ρ_mm_*), (e)(f) Effect of the fixed cost of choosiness (*c_f_*), (g)(h) Effect of the relative cost of choosiness (*c_r_*). The default parameters values are as follows: *ρ_MM_* = 0, *ρ_mm_* = 0.01, *h_a_* = 1, *h_m_* = −1, *r* = 0.1, *c_r_* = 0, *C_f_* = 0, *S_aa_* = 0, *S_ab_* = 0.01 and *S_bb_* = 0.

